# Hypermutation in *Cryptococcus* reveals a novel pathway to 5-fluorocytosine (5FC) resistance

**DOI:** 10.1101/636928

**Authors:** R. Blake Billmyre, Shelly Applen Clancey, Lucy X. Li, Tamara L. Doering, Joseph Heitman

**Author notes:** These authors contributed equally to this work (R. Blake Billmyre, Shelly Applen Clancey). Stowers Institute for Medical Research, 1000 E 50^th^ St., Kansas City, MO 64110, USA. Corresponding author: Room 322 CARL Building, Box 3546 Research Drive, Department of Molecular Genetics and Microbiology, Duke University Medical Center, Durham, NC 27710, USA.

## Abstract

Drug resistance is a critical challenge in treating infectious disease. For fungal infections, this issue is exacerbated by the limited number of available and effective antifungal agents. Patients infected with the fungal pathogen *Cryptococcus* are most effectively treated with a combination of amphotericin B and 5-fluorocytosine (5FC). Isolates causing infections frequently develop resistance to 5FC although the mechanism of this resistance is poorly understood. Here we show that resistance is acquired more frequently in isolates with defects in DNA mismatch repair that confer an elevated mutation rate. Natural isolates of *Cryptococcus* with mismatch repair defects have recently been described and defective mismatch repair has been reported in other pathogenic fungi. In addition, whole genome sequencing was utilized to identify mutations associated with 5FC resistance *in vitro*. Using a combination of candidate-based Sanger and whole genome Illumina sequencing, the presumptive genetic basis of resistance in 16 independent isolates was identified, including mutations in the known resistance genes *FUR1* and *FCY2*, as well as a novel gene, *UXS1*. Mutations in *UXS1* lead to accumulation of a metabolic intermediate that appears to suppress toxicity of both 5FC and its toxic derivative 5FU. Interestingly, while a *UXS1* ortholog has not been identified in other fungi like *Saccharomyces cerevisiae*, where the mechanisms underlying 5FC and 5FU resistance were elucidated, a *UXS1* ortholog is found in humans, suggesting that mutations in *UXS1* in cancer cells may also play a role in resistance to 5FU when used during cancer chemotherapy in humans.

## Introduction

One of the key challenges of the 21^st^ century is the emergence and reemergence of pathogens. Opportunistic fungal pathogens comprise an important component of this problem as they infect the rapidly expanding cohort of immunocompromised patients [1]. These pathogens are responsible for millions of infections annually, with substantial mortality. Among the most dangerous are *Cryptococcus* species that cause approximately 220,000 infections a year, with more than 181,000 attributable deaths [2]. Cryptococcosis is particularly prominent in Sub-Saharan Africa, where the HIV/AIDS epidemic has resulted in a large population of susceptible individuals. Cryptococcosis is treated most effectively using a combination of 5-fluorocytosine (5FC) and amphotericin B [3,4]. However, in the parts of Africa where patients are most commonly afflicted with cryptococcosis, the medical infrastructure is insufficient to allow treatment with the highly toxic amphotericin B component of this dual therapy. Instead patients are typically treated with fluconazole monotherapy, with limited success. Excitingly, recent studies have shown that 5FC can be effectively paired with fluconazole to replace amphotericin B for treatment of patients in Africa [5]. However, 5FC is not yet approved or available for treatment in any African countries.

5FC acts as a prodrug, which enters cells via the cytosine permease Fcy2. 5FC itself is not toxic, but upon uptake into fungal cells, it is converted into toxic 5-fluorouracil (5FU) by cytosine deaminase, an enzyme that is not present in human cells [6]. In *Cryptococcus*, and other fungi, cytosine deaminase is encoded by the *FCY1* gene. 5FU is then further processed by the product of the *FUR1* gene, a uracil phosphoribosyltransferase, and inhibits both DNA and protein synthesis. Resistance is well understood in other fungal pathogens, like *Candida albicans*, where loss of function mutations in *FCY1, FCY2*, and *FUR1* can mediate resistance to 5FC [7]. In *Candida lusitaniae*, mutations in *FUR1* can be readily distinguished from mutations in *FCY1* and *FCY2* because only *fur1* mutations result in cross-resistance to 5FU [8]. Likewise, in *Candida dubliniensis*, natural missense *fur1* mutations affect both 5FC and 5FU resistance [9]. However, little work has been conducted on 5FC resistance directly in *Cryptococcus*. One of the few early studies suggested that reductions in *FUR1* activity may be linked to resistance to 5FC based on a high frequency of cross-resistance to 5FU [10]. However, this study took place prior to the cloning or sequencing of the *FUR1* gene in *Cryptococcus* and attribution of resistance to *FUR1* was based only on cross-resistance to 5FU. More recent studies of 5FC resistant *Cryptococcus bacillisporus* isolates found no mutations in *FCY1, FUR1*, or any of three putative *FCY2* paralogs that explained drug resistance [11]. However, in *Cryptococcus deuterogattii*, deletions of *FCY2* confer resistance to 5FC [12].

Recent work has demonstrated one source of increased rates of resistance to antifungal drugs in *Cryptococcus*: defects in the DNA mismatch repair pathway [13,14]. Natural isolates with DNA mismatch repair defects have been identified in both an outbreak population of *Cryptococcus deuterogattii* [13,15] and in *Cryptococcus neoformans* [14,16]. Defects in mismatch repair are also common in other human fungal pathogens, including *Candida glabrata* [17]. Depending on the population studied, multidrug resistance is sometimes linked to the hypermutator state in *C. glabrata* [18,19]. A recent study of clinical *C. glabrata* isolates in India found a high prevalence of *msh2* mutation, but no drug resistance [20]. This could suggest that hypermutation is advantageous even prior to drug exposure, while also providing more rapid development of resistance when antifungal drugs or agents are encountered. Alternatively, hypermutation has also been observed in more ancient lineages of fungi not known to be pathogenic, suggesting that hypermutation may have general advantages in a broader range of settings [21]. Here we demonstrate that DNA mismatch repair defects also enable rapid resistance to 5FC in *C. deuterogattii* (previously known as *C. gattii* VGII [22–24]). We then utilize whole genome Illumina sequencing, in combination with candidate-based Sanger sequencing, to identify the genetic basis for drug resistance in 16 independent isolates. We attribute resistance to mutations in *FUR1* and unexpectedly, we also identify a novel pathway of resistance to 5FC involving mutations in the pathway responsible for producing the capsule, a core component of cryptococcal virulence.

## Results

In a previous study, we demonstrated that mismatch repair mutations conferred increased rates of resistance to the antifungal drugs FK506 and rapamycin [13]. Because these hypermutator strains are found among both environmental and clinical isolates, here we tested if a hypermutator state could also confer resistance to one of the front-line drugs used to treat Cryptococcosis: 5-fluorocytosine (5FC). A semi-quantitative swabbing assay was first employed to demonstrate that deletions of the mismatch repair gene *MSH2* in *Cryptococcus deuterogattii* confer an elevated rate of resistance to 5FC (Figure 1A). This result was confirmed using a quantitative fluctuation assay approach (Figure 1B). This assay revealed a greater than 15-fold increase in the generation of resistance to 5FC in *msh2*Δ mismatch repair defective mutants. Similarly, a simple spreading assay using VGIIa-like strains that had previously been found to harbor an *msh2* nonsense allele [13] demonstrated a much higher rate of resistance to both 5FC and 5FU than in the VGIIa non-hypermutator strains (Supplemental Figure 1).

**Figure 1.**
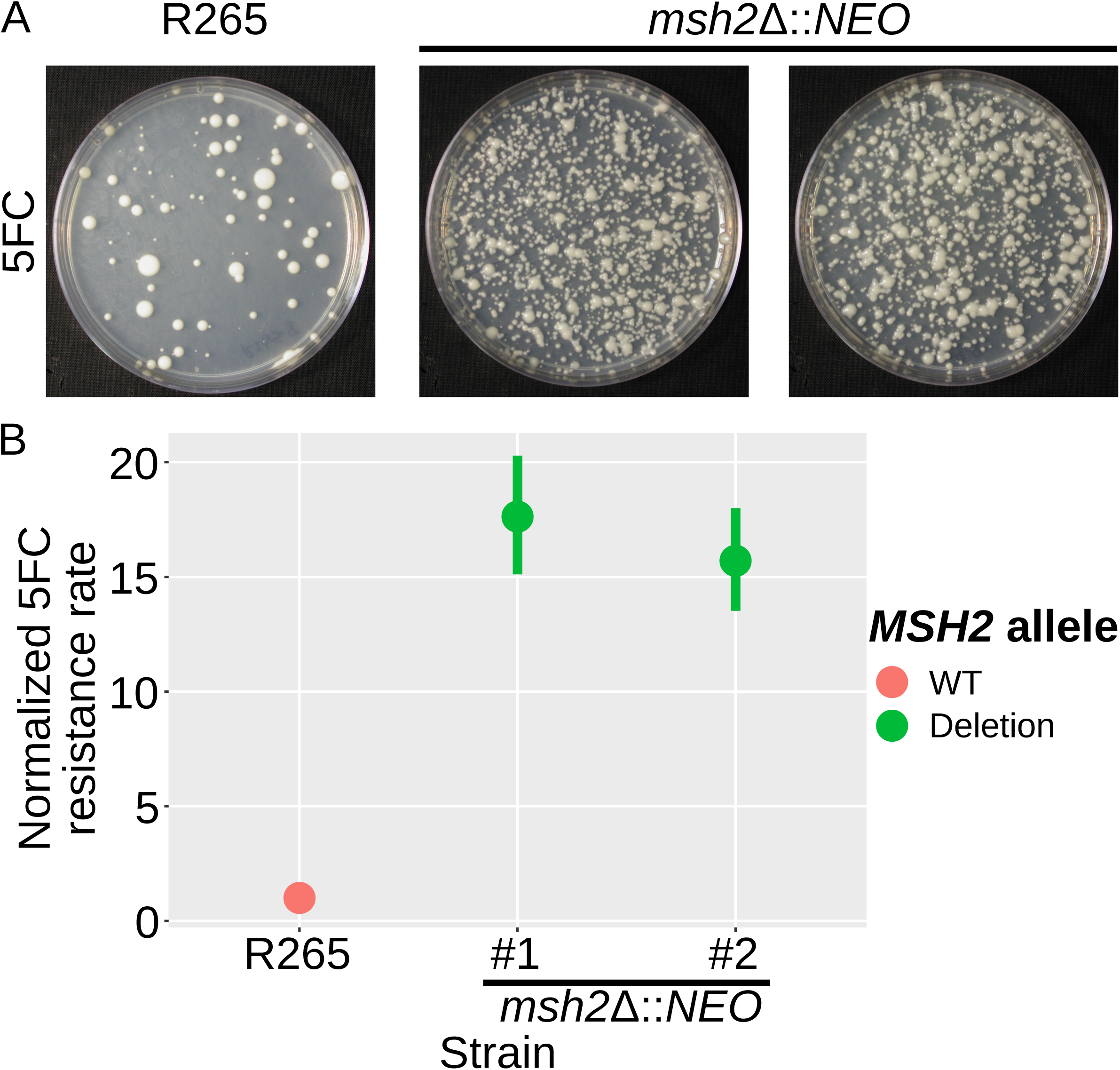
5FC resistance is enhanced by defects in mismatch repair. A) Swab assays were conducted using both the wildtype R265 strain and two independent *msh2*Δ∷*NEO* mutants to test for the ability to generate resistance to 5FC. All three strains developed resistance; however, the mismatch repair mutants generated resistant isolates at a higher frequency. B) A fluctuation assay was conducted to compare 5FC resistance quantitatively between wildtype R265 and two independent *msh2*Δ∷*NEO* mutants. Mutation rate was normalized to the wildtype strain. Both mutator strains showed a greater than 15-fold increase in the rate of resistance.

In previous studies, mutator alleles in *C. deuterogattii* were not found to be generally advantageous in rich media [13]. However, under stressful conditions, such as drug challenge with FK506 and rapamycin, mutator alleles were highly beneficial. A competitive growth experiment was utilized to test the same concept with 5FC. Mutator strains became resistant to 5FC at a higher rate and thus rapidly outcompeted wildtype strains (Figure 2). However, in the absence of added stress, the mutator alleles showed no such advantage. This result suggests that drug challenge during infection may select for strains with elevated mutation rates that are able to acquire drug resistance more rapidly.

**Figure 2.**
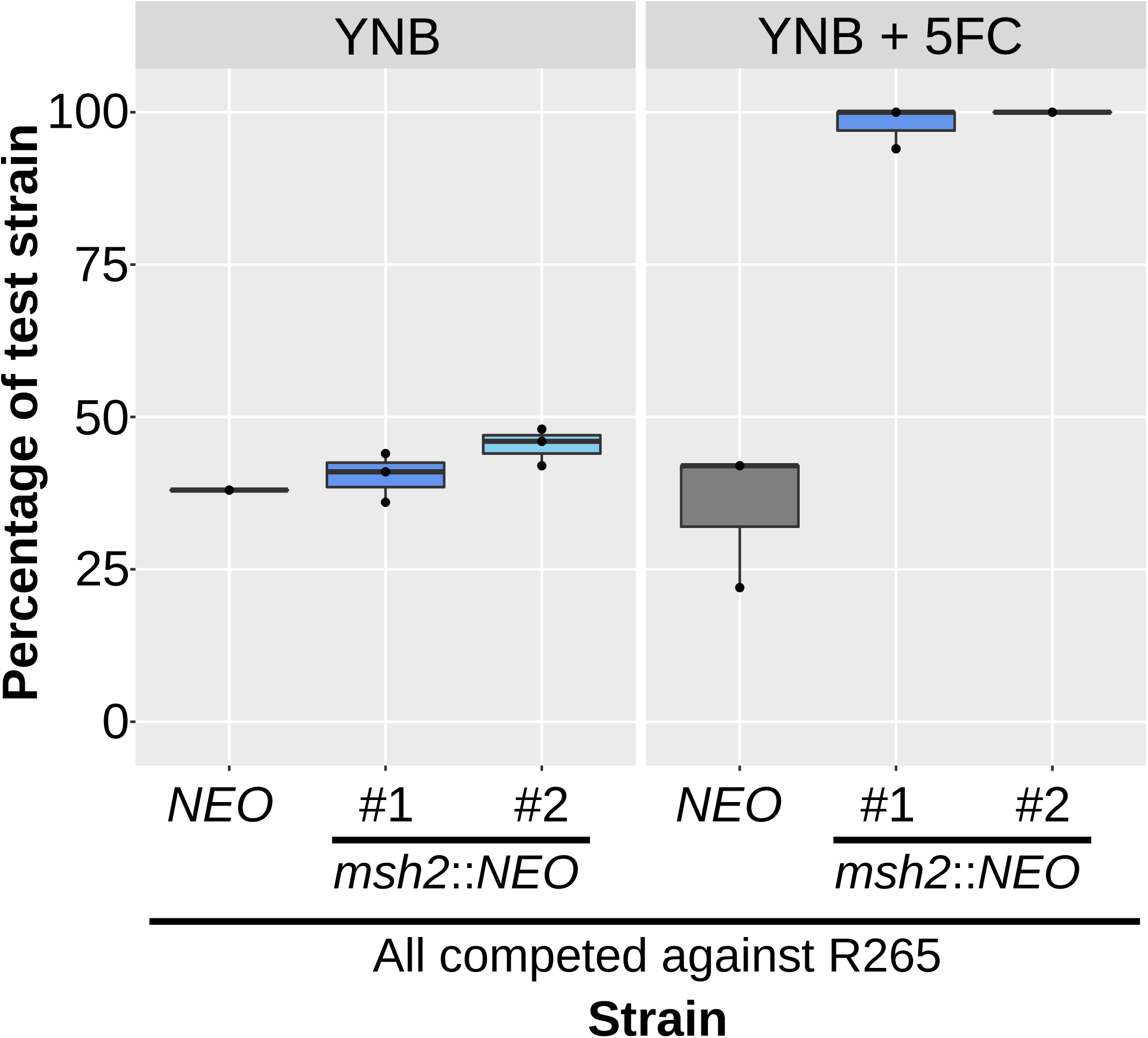
Exposure to 5FC generates an adaptive advantage for mutator strains. Competition experiments between a tester strain with a neomycin resistance marker and a wildtype R265 strain. (Strain used: SEC501, RBB17, RBB18). Overnight cultures were mixed 1:1 and then used to inoculate a second overnight culture in liquid YNB with and without 5FC. All three marked strains showed a slight growth defect in comparison to the unmarked strain in nonselective media but only the hypermutator strains demonstrated a dramatic growth advantage when grown in YNB+5FC. Boxplots show minimum, first quartile, median, third quartile, and maximum values. Points represent the results from three individual replicates and are summarized by the box plot. The R265 *NEO*^R^ vs wildtype competition is gray, while thetwo *msh2*Δ∷*NEO* vs wildtype competitions are dark and light blue.

In other fungi, resistance to 5FC is typically mediated by mutations in one of three genes: *FCY1, FCY2*, or *FUR1* [7,8,10,25]. As described above, mutations in *FCY1* and *FCY2* are typically distinguishable from *fur1* mutations because mutations in *FUR1* confer resistance not only to 5FC but also to 5FU. In contrast, *fcy1* and *fcy2* mutations confer resistance to only 5FC. To define the mechanism underlying 5FC resistance in *C. deuterogattii*, 29 resistant colonies were isolated and tested, originating from the wildtype (R265, 9 colonies) and from two independent *msh2*Δ mutants derived in the R265 background (RBB17, 10 colonies and RBB18, 10 colonies). Cultures were started from independent colonies and a single resistant colony was selected from each culture, so that only one resistant isolate is derived from any original colony derived from the frozen stock. All of the 5FC resistant isolates (Table 1) acquired were cross-resistant to 5FU (29/29) (Figure 3A), leading us to hypothesize that resistance to 5FC in *C. deuterogattii* was most commonly mediated by mutations in *FUR1.*

**Table 1.** 5FC-resistant isolates whole genome sequenced or successfully genotyped by Sanger sequencing.

| Strain Name | Original Genotype | Putative Resistance Allele |
| --- | --- | --- |
| R265-1 | Wildtype | *fur1* 1134delT |
| R265-2 | Wildtype | ~18.5 kb deletion spanning *fur1* |
| R265-3 | Wildtype | *fur1* 1003delT, mutation detected via Sanger |
| R265-4 | Wildtype | *fur1* 1136delT, mutation detected via Sanger |
| R265-5 | Wildtype | *uxs1* 1520delT |
| R265-6 | Wildtype | *fur1* 1440delA, mutation detected via Sanger |
| R265-7 | Wildtype | ~14.7 kb deletion spanning *fur1* |
| R265-8 | Wildtype | ~26.7 kb deletion spanning *fur1* |
| RBB17-1 | *msh2Δ::NEO* |  |
| RBB17-2 | *msh2Δ::NEO* |  |
| RBB17-3 | *msh2Δ::NEO* |  |
| RBB17-4 | *msh2Δ::NEO* |  |
| RBB17-5 | *msh2Δ::NEO* | *fur1* 1358delA in 6 base homopolymer |
| RBB17-6 | *msh2Δ::NEO* |  |
| RBB17-7 | *msh2Δ::NEO* |  |
| RBB17-8 | *msh2Δ::NEO* | *fur1* G448A (splice acceptor) |
| RBB18-1 | *msh2Δ::NEO* |  |
| RBB18-2 | *msh2Δ::NEO* | *fur1* 1358delA in 6 base homopolymer |
| RBB18-3 | *msh2Δ::NEO* |  |
| RBB18-4 | *msh2Δ::NEO* | *fcy2* Trp167Stop<br>*uxs1* Asp306Gly |
| RBB18-5 | *msh2Δ::NEO* | *fur1* 1358delA in 6 base homopolymer |
| RBB18-6 | *msh2Δ::NEO* | *fur1* 1027delT in 5 base homopolymer |
| RBB18-7 | *msh2Δ::NEO* |  |
| RBB18-8 | *msh2Δ::NEO* | *uxs1* 1182insC in 7 base homopolymer |
| RBB18-9 | *msh2Δ::NEO* | *uxs1* Tyr217Cys |

**Figure 3.**
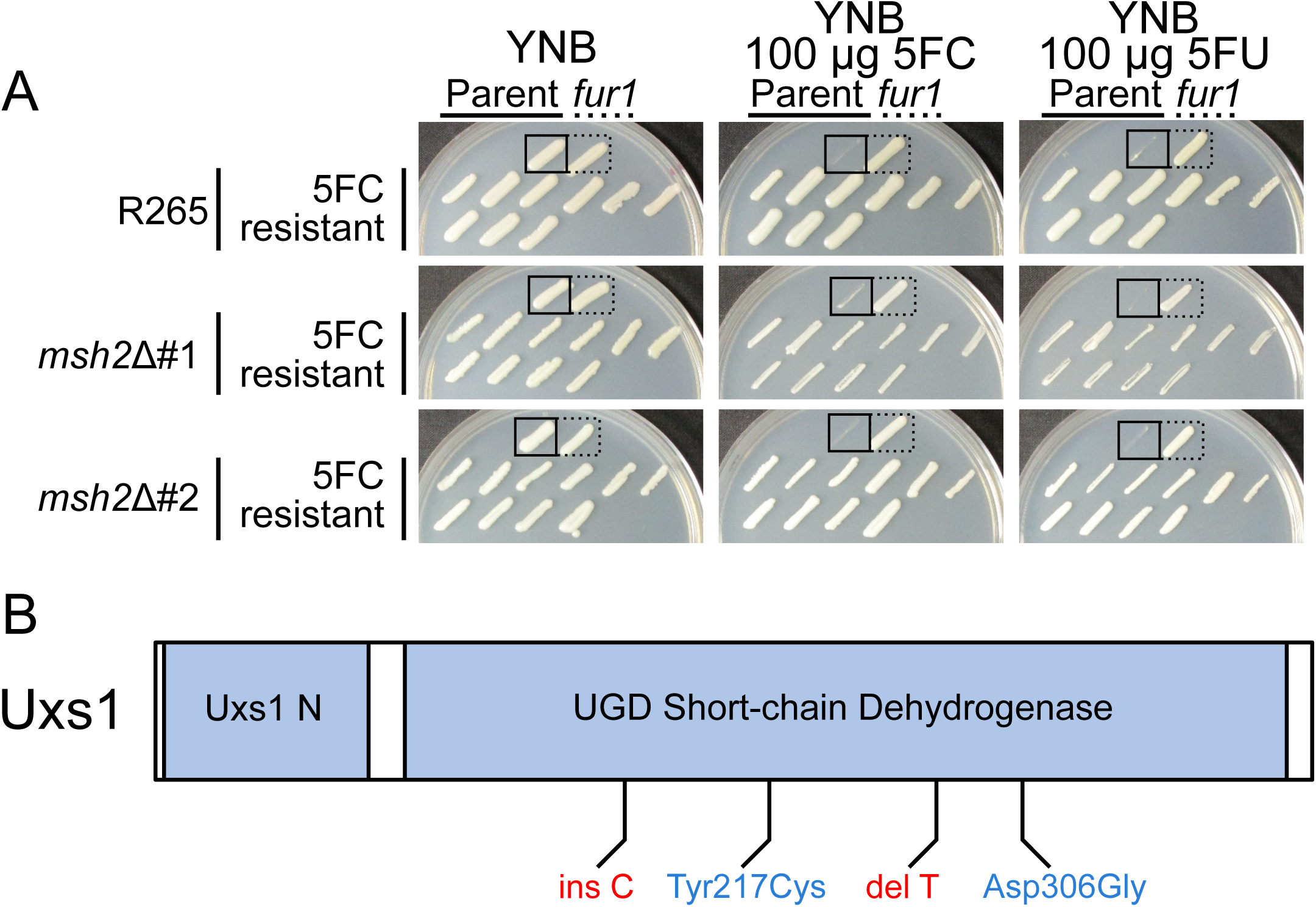
5FC resistant mutants are cross-resistant to 5FU. A) Isolates that were selected based on growth on 5FC media were patched to YNB, YNB plus 5FC, and YNB plus 5FU. Each plate has parental and *fur1* mutant controls in the top row. Hypermutator controls have occasional resistant colonies that emerged in the growth patch. Sanger sequencing revealed that very few isolates had sustained mutations in *FUR1*. B) Schematic showing the predicted domains encoded by the *UXS1* gene as well as the location and number of mutations identified. Nonsense alleles are shown in red and missense are shown in blue.

However, when the *FUR1* gene was sequenced in this set of 5FC/5FU resistant isolates, unexpectedly, only three out of 29 isolates (10.3%) were found to have sustained mutations in *FUR1* (R265-3, R265-4, and R265-6) (Table 1), although PCR amplification of the *FUR1* locus failed for another 3 isolates (R265-2, R265-7, R265-8), suggesting a possible large deletion or insertion event. Because *fur1* mutations were the only known cause of 5FC/5FU cross-resistance, we performed whole genome Illumina sequencing on a subset of the remaining isolates (22 isolates) to identify unknown genes underlying resistance. We sequenced the whole genomes of 5 additional R265 isolates, 8 additional RBB17 isolates, and 9 additional RBB18 isolates, for a total of 22 5FC and 5FU resistant isolates.

From the sequenced genomes, reads were aligned to the R265 reference genome and SNPs and indels were identified. This analysis revealed that one pair of the presumed independent isolates were in fact siblings (RBB17-3 and RBB17-4), resulting in a total of 5 independent R265 genomes, 7 independent RBB17 genomes, and 9 independent RBB18 genomes (21 total independent isolates).

Of these 21 independent genome sequences, six contained unambiguous mutations in *FUR1* that were not detected by Sanger sequencing. The first *fur1* mutation discovered by whole genome sequencing was a single base deletion that introduced a frameshift (R265-1). Two sets of homopolymer shifts were also identified in *FUR1*: a single base deletion in a 6xA homopolymer run at position 1358 found in three independent isolates (RBB17-5, RBB18-2, and RBB18-5) and a single base deletion in a 5xT homopolymer run at position 1027. Finally, a mutation within a splice acceptor (G to A) was identified at position 448 (RBB17-8).

For three 5FC resistant R265 strains (R265-2, R265-7, R265-8), PCR amplification of the *FUR1* locus failed and subsequent whole genome sequencing revealed regional deletions consistent with these failed PCRs. For two strains, break points were clearly identifiable. R265-7 sustained a deletion of bases 189022-203758 (14,736 bp) surrounding *FUR1*, while R265-8 sustained a deletion of bases 190136-216860 (26,724 bp), also including *FUR1*. For R265-2, one end of the deletion lies within *FUR1*, eliminating one of the primer binding sites and consistent with the failed PCR. The other end of the ∼18.5 kb deletion fell within a sequencing gap of the annotated V2 R265 reference genome. To identify the precise location of this second breakpoint, reads from R265-2 were mapped to a recent Nanopore and Illumina hybrid assembly of the R265 strain [26]. Interestingly, the second breakpoint was found within a gene encoding a weak paralog of *FUR1* (5 × 10^−10^ protein BLAST e-value). This paralog (CNBG_4055) is also present in *C. neoformans* (CNAG_2344), suggesting that if it arose via duplication, it was before the last common ancestor to both species. Given that deletion of *FUR1* confers resistance to 5FC and 5FU, it is unlikely that this paralog performs the same function as Fur1 (Figure 3A). Despite the protein similarity, no obvious nucleotide homology was found that may have mediated this large deletion conferring 5FC resistance. In fact, the *FUR1* paralog is inverted relative to *FUR1*, reducing the likelihood that remnant homology may have generated a region susceptible to frequent homology-mediated deletion of *FUR1* that would yield the type of regional deletion observed here.

A Trp167STOP mutation in *FCY2* (CNBG_3227) was also detected in the sequenced set (RBB18-4). Mutations in *FCY2* were unexpected because in other fungi they do not confer resistance to 5FU and because there are 2 additional paralogs with substantial similarity to *FCY2* present in the *Cryptococcus* genome. Because this *fcy2* strain also contains a second mutation in a gene that plays a role in 5FC and 5FU resistance (discussed below), the *fcy2* mutation may be unrelated to drug resistance or may enhance resistance in the presence of the second mutation. We attempted to test the ortholog of *FCY2* from *Cryptococcus neoformans* using a deletion collection strain but found that the mutant in the collection retained a functional copy of the *FCY2* gene. However, *fcy2* deletion has recently been reported to confer resistance to 5FC in *C. deuterogattii* [12].

In total, out of 29 original 5FC resistant strains (Table 1), twelve independent *fur1* mutations were identified using Sanger and Illumina sequencing. One independent *fcy2* mutation was identified by Illumina sequencing. We did not identify any *fcy1* mutations, although *fcy1* mutations confer resistance to 5FC in *C. neoformans* (Supplemental Figure 2). In total, 11 sequenced genomes representing 10 independent isolates remained with no mutations in any genes previously described to have a role in 5FC or 5FU resistance. These genomes were examined to identify novel candidate mutations. To distinguish causal variants from background mutations, candidate genes were required to be mutated in at least two different independent isolates. Variant impact was also scored using SNPeff [27] and mutations were not considered if predicted to have low impact (i.e., synonymous, intronic, or non-coding variants). Mutations of moderate or higher impact were identified at a total of 128 sites (Supplemental Table 3). To further prioritize, we specifically focused on mutations that were present in isolates from more than one of the parental backgrounds. We identified *UXS1*, which sustained four novel mutations in four isolates from two parental backgrounds (Figure 3B).

*UXS1* encodes the enzyme that converts UDP-glucuronic acid to UDP-xylose [28]. This pathway is critical for the formation of the capsule, a core virulence trait of *Cryptococcus*, and for synthesis of other glycoconjugates. There is no *UXS1* ortholog in either *Saccharomyces cerevisiae* or *Candida albicans*, where many of the resistance mechanisms for 5FC were elucidated. The mutations in *UXS1* included a single base deletion in a 3xT homopolymer (R265-5), a single base insertion in a 7xC homopolymer (RBB18-8), and a missense mutation (Tyr217Cys, RBB18-9) (Figure 3B, Table 1). Finally, a *uxs1* mutation (Asp306Gly) was identified in the isolate previously identified to have an *fcy2* mutation (RBB18-4). In sum, 9 sequenced genomes representing 8 independent isolates remained for which we were unable to identify a mutation that conferred resistance to 5FC and 5FU, all derived from *msh2* mutant isolates.

To confirm the role of *uxs1* mutation in resistance to 5FC and 5FU, a *uxs1* deletion available from a *C. neoformans* deletion collection was employed (Figure 4A). This *uxs1*Δ strain was completely resistant to both drugs, suggesting that all three alleles isolated were likely loss of function mutations because they shared a drug resistance phenotype with the null mutant. We tested the MIC of 5FC for *uxs1* and *fur1* mutants in both YPD and YNB using a broth microdilution assay. Both *uxs1* and *fur1* mutants were resistant to 5FC above the limits of our assays (MIC > 400 µg/mL in YPD and >4 µg/mL in YNB) while the wildtype parent strains were sensitive at 200 µg/mL in YPD and 0.5 µg/mL in YNB (Table 2).

**Table 2.**
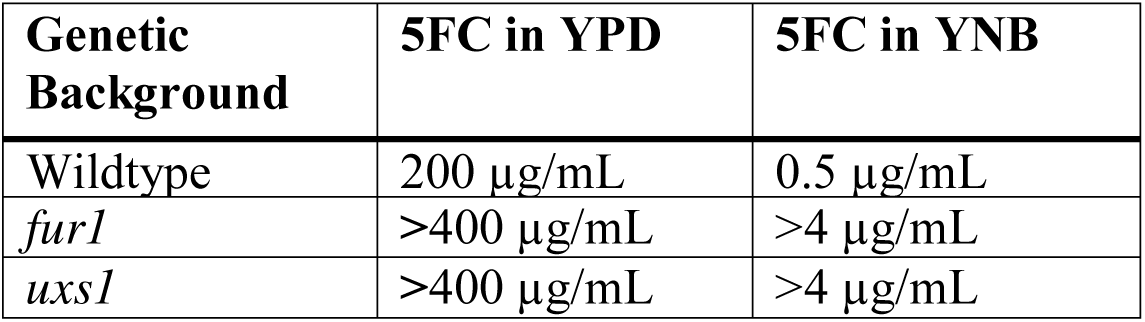
MIC values for deletion mutants of genes identified in this study.

**Figure 4.**
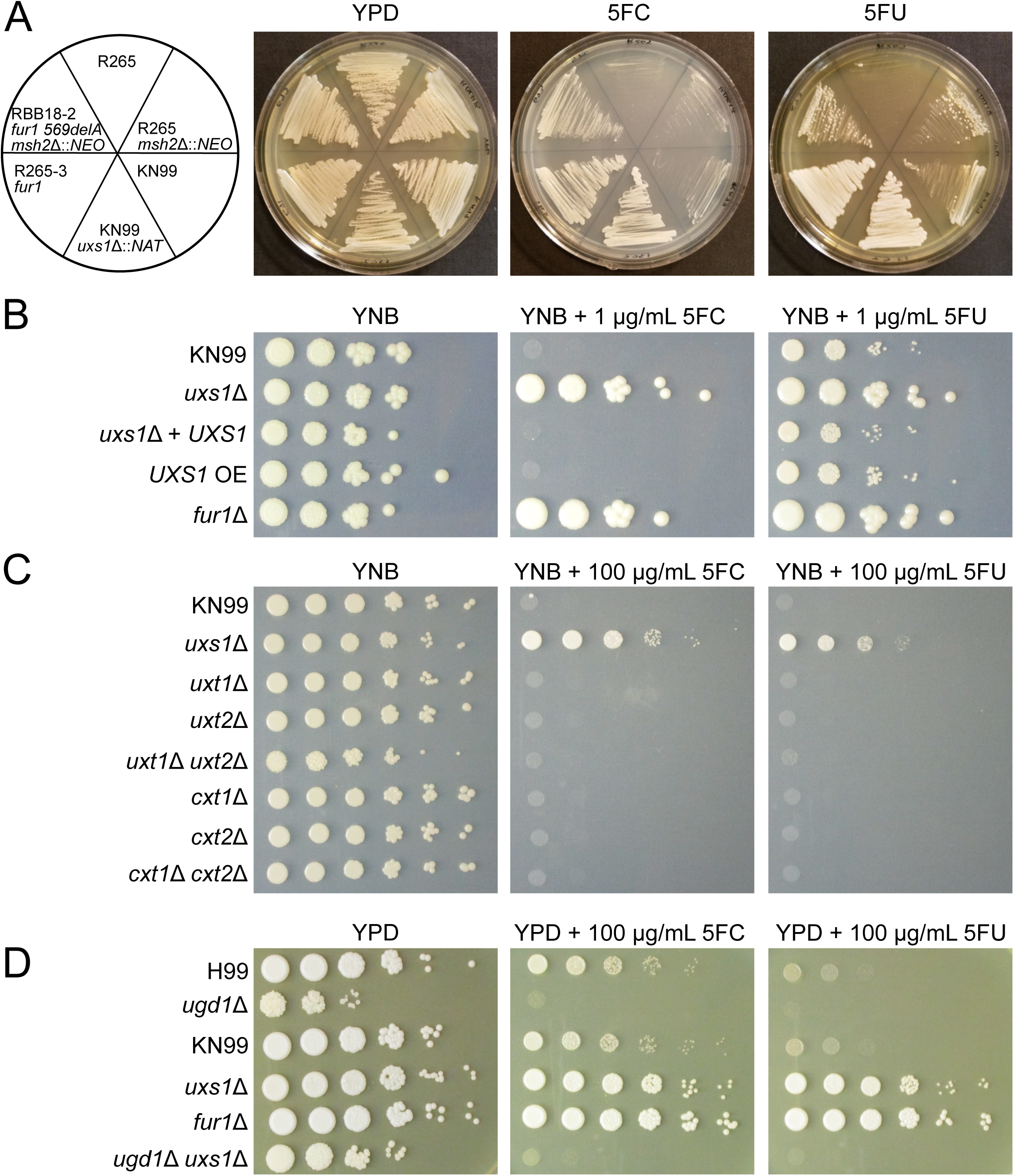
*uxs1* mutants mediate 5FC resistance through a xylosylation-independent mechanism. A) KN99 deletion strains from the *C. neoformans* deletion collection show that deletion of *UXS1* confers resistance to 5FC and 5FU. The RBB18-2 strain carrying a *fur1* mutation is resistant to 5FC and 5FU although more weakly to 5FU. The R265-3 strain carrying a *fur1* mutation is completely resistant to both drugs. B) Spot dilution assay on YNB, YNB plus 5FC, and YNB plus 5FU demonstrating overexpression of *UXS1* driven by the actin promoter does not confer increased sensitivity to 5FC or 5FU. C) Spot dilution assays on YNB, YNB plus 5FC, and YNB plus 5FU demonstrating that mutants deficient in UDP-xylose transport (*uxt1*Δ, *uxt2*Δ, *uxt1*Δ *uxt2*Δ) and xylose transferase mutants (*cxt1*Δ, *cxt2*Δ, *cxt1*Δ *cxt2*Δ) show no change in 5FC and 5FU sensitivity. D) Spot dilution assay on YPD, YPD plus 5FC, and YPD plus 5FU showing that *ugd1* mutants are viable on rich YPD media but retain sensitivity to 5FC and 5FU. In addition, *ugd1 uxs1* double mutants retain sensitivity to 5FC and 5FU like a *ugd1* single mutant rather than gain resistance like the *uxs1* single mutant.

We next sought to genetically define the mechanism by which drug resistance may be mediated by loss of *uxs1* function. Multiple models were considered to explain why 5FC/5FU toxicity would require Uxs1. The first was that Uxs1 directly converts 5FU into a toxic product. If so, Uxs1 and Fur1 would function in the same pathway, as either mutant independently confers drug resistance. This hypothesis was tested using an overexpression allele of *UXS1* that is driven by the actin promoter [29]. If this hypothesis were correct, we would expect to observe additional sensitivity conferred by the overexpression allele compared to wildtype. By reducing the amount of 5FU used to only 1 µg/mL, wildtype strains were only partially inhibited. However, introduction of an overexpression allele of *UXS1* did not increase sensitivity (Figure 4B). This suggests that Uxs1 does not act by converting 5FU or a 5FU derivative into a toxic product.

We next tested whether 5FC resistance in *uxs1* mutants may occur through an indirect effect of the role of Uxs1 in synthesis of UDP-xylose. UDP-xylose is the donor molecule for xylose addition to glycans, a process that primarily occurs in the secretory compartment. If xylosylation of an unknown glycoconjugate is required to mediate 5FC toxicity, mutation of *UXS1* would indirectly confer drug resistance. To test this, deletion mutants lacking transporters that move UDP-xylose into the secretory compartment (*uxt1, uxt2*, and a *uxt1 uxt2* double mutant [30]) or that lack Golgi xylosyl-transferases that act in protein, glycolipid, and polysaccharide synthesis (*cxt1* [31], *cxt2*, and a *cxt1 cxt2* double mutant) were analyzed. None of these mutants demonstrated any change in sensitivity to 5FC or 5FU (Figure 4C). However, these data did not rule out a requirement for a (previously undescribed) cytoplasmic xylosyl protein modification. To test this hypothesis, a mutant that cannot generate UDP-glucuronic acid, the immediate precursor for UDP-xylose synthesis was used. This mutant (*ugd1*) is somewhat growth impaired relative to wildtype and cannot grow on YNB media. However, it does grow, albeit poorly, on rich YPD media, where it clearly exhibited sensitivity to 5FC. This result demonstrated that xylose modification, in any cellular compartment, is not required for 5FC toxicity (Figure 4D).

The previous models ruled out the lack of UDP-xylose for synthetic processes as an explanation for 5FC resistance. Another result of the loss of *UXS1* function is the accumulation of UDP-glucuronic acid, the immediate precursor in the production of UDP-xylose. Past studies have shown that UDP-glucuronic acid accumulates to extremely high levels in *uxs1* mutant cells, while it is undetectable in *ugd1* mutants [32]. To test whether this mediates resistance, we generated a *uxs1 ugd1* double mutant, which should produce neither UDP-glucuronic acid nor UDP-xylose [32]. While the *uxs1 ugd1* mutant was growth impaired, like the *ugd1* single mutant, it was clearly sensitive to 5FC (Figure 4D). That *uxs1* mutants are 5FC resistant, whereas *uxs1 ugd1* double mutants are restored to 5FC sensitivity suggests that accumulation of UDP-glucuronic acid in *uxs1* mutants mediates resistance to 5FC and 5FU (Figure 5).

**Figure 5.**
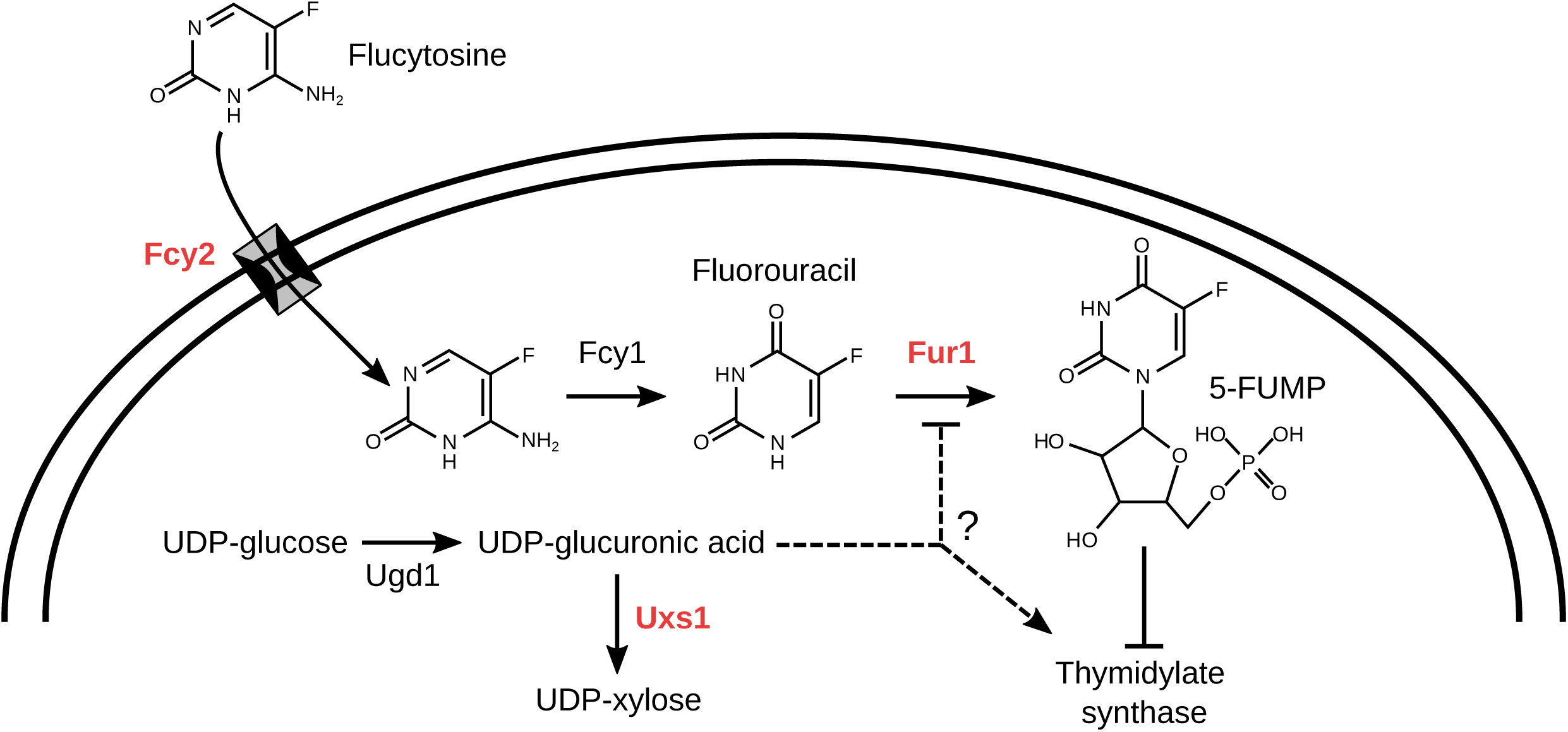
Model of inhibition of 5FC/5FU toxicity by *uxs1* mutation. Potential mechanisms by which *uxs1* mutations may confer resistance to both 5FC and 5FU. Mutation of *uxs1* causes an accumulation of UDP-glucuronic acid, the product of Ugd1, which either impairs production of toxic fluoridated molecules or rescues inhibition of the targets of those fluoridated molecules, such as thymidylate synthase. Protein names are in red for those where mutations were found in this study.

## Discussion

Treating fungal diseases is complicated both by the limited number of drugs that effectively treat infection without harming the patient and by the rapid rate at which fungi develop resistance to the few drugs that are effective. 5FC is a particularly emblematic example of this issue, as it is highly efficacious with limited toxicity. Human cells lack the ability to convert 5FC to 5FU and toxicity is conferred only by the conversion of 5FC to the chemotherapeutic 5FU by a patient’s microbiota [33]. However, 5FC is ineffective when used for solo treatment because fungal resistance rapidly emerges. Here, we demonstrate that DNA mismatch repair mutants exhibit accelerated acquisition of resistance to 5FC. Evolutionary theory predicts that hypermutators should be rare in eukaryotic microbes because sex unlinks mutator alleles from the mutations they generate, eliminating the advantage of an elevated mutation rate and leaving only the general decrease in fitness from introduced mutations [34]. This result lends further support to the recent appreciation that mismatch repair mutants may be common in pathogenic fungi in part because treatment with antifungal drugs increases selection for mutations that generate resistance [13,14,16,17].

We explored the underlying genetic and genomic basis of 5FC resistance. The resistant mutants in *C. deuterogattii* selected here were cross-resistant to 5FU. Sanger and whole genome Illumina sequencing identified a presumptive genetic basis for drug resistance in 16 independent isolates. Analysis of resistance loci from whole genome data was relatively facile in wildtype strains (5/5 strains assigned a causative mutation), where an average of 1.2 coding mutations (range 0-3) were identified by whole genome sequencing, including the putative resistance mutation, relative to the reference. However, this analysis was substantially more difficult in mutator strains (8/17) where an average of 11.47 coding mutations were found per strain (range 2-25), with numerous additional noncoding or synonymous mutations. For the purposes of identifying the genetic basis of a trait that occurs at a high rate in wildtype, future studies would be advised to avoid mutations that increase mutation rate, as they contribute to background noise.

We identified multiple mutations in the *FUR1* locus (12 of the 16 identified causative mutations). *fur1* mutations occurred through multiple mechanisms, including regional deletions, homopolymer tract length changes that introduced frameshift mutations, and a splice site acceptor point mutation. Surprisingly, we did not identify mutations in *FCY1* or mutations in *FCY2* that were unaccompanied by a second resistance mutation. Although we selected with only 5FC, all drug resistant isolates were cross-resistant to 5FU as well. One possible explanation is that selection with 100 µg/mL of 5FC may be above the MIC for *fcy1* or *fcy2* mutants in *C. deuterogattii*, although *fcy1* mutants in *C. neoformans* are resistant to 100 µg/mL of 5FC (Supplemental Figure 2). Further experiments will be necessary to test this hypothesis, which could provide guidance into treatment levels for 5FC. Further experiments based on this hypothesis could provide insight into the function of the other *FCY2* paralogs, perhaps as lower affinity transporters of 5FC that confer toxicity at higher concentrations of 5FC.

Mutations in *UXS1* are particularly interesting as a mechanism of resistance in *Cryptococcus* because Uxs1 catalyzes the production of UDP-xylose, the donor molecule for essential components of Cryptococcal capsule polysaccharides. Strains lacking *UXS1* are hypocapsular with altered capsule structure [32]. In addition, *uxs1* mutants are avirulent in a murine tail-vein injection disseminated infection model [35]. This suggests that *uxs1* mutants might be unlikely to emerge during exposure to 5FC *in vivo*, even though they represent a substantial proportion of the resistant isolates observed in this study. Likewise, regional deletions including *FUR1* affected multiple neighboring genes as well, including the direct neighboring gene *GIS2*. Gis2 has previously been described to play a role in stress tolerance, including fluconazole and oxidative stress tolerance [36]. Like *uxs1* mutants, these regional deletion mutants may be less likely to emerge *in vivo*. It is important to note that *in vitro* resistance to 5FC is not necessarily associated with clinical treatment failure and does not prevent synergy of combination treatment with Amphotericin B and flucytosine [37]. Continued selection by 5FC treatment of a deleterious resistance allele like a *uxs1* mutation or a collateral *gis2* deletion might explain the maintenance of synergy. Future studies examining the mechanisms of resistance during treatment with 5FC *in vivo* will provide further insights into the possible contribution of each of these mechanisms to resistance in patients.

This study also illustrates the importance of examining drug resistance in the context of the pathogen being treated. Previous work in *C. albicans* and *S. cerevisiae* suggested that resistance would occur through mutations in *FUR1*, but both species are evolutionarily distant from *Cryptococcus* and lack a *UXS1* ortholog. While these previous studies provided substantial insight into 5FC toxicity, studies in the pathogen of interest are essential. Surprisingly, one strain (RBB18-4) that was cross resistant to 5FU had a mutation in the *FCY2* gene (CNBG_3227), which in other species confers resistance to 5FC but not 5FU. Mutation of *FCY2* is known to result in resistance to 5FC in *C. deuterogattii*, but cross-resistance to 5FU has not been tested [12]. Unexpected cross-resistance between 5FC and fluconazole has been previously observed in *fcy2* mutants of *Candida lusitaniae* but is proposed to occur through competitive inhibition of fluconazole uptake by 5FC that can no longer enter through Fcy2-mediated transport [8,38,39]. *C. lusitaniae fcy2* mutants are not resistant to fluconazole without the addition of 5FC. In addition, multiple resistant strains were not assigned a presumptive causative mutation here and lacked mutations in any genes known to cause 5FC resistance from this or previous work (*FUR1, FCY1, FCY2*, and *UXS1*). Presumably unknown mechanisms are responsible for resistance to 5FC and 5FU in these strains as well, either in pathways unique to *Cryptococcus* or potentially more broadly conserved.

In addition, *UXS1* mutations provide unexpected insight into interaction between nucleotide synthesis and generation of precursors for xylosylation. Surprisingly, accumulation of UDP-glucuronic acid appears to either inhibit the pyrimidine salvage pathway or activate thymidylate synthase (Figure 5). This suggests that UDP-glucuronic acid may have a role as a source of UDP for the cell, while UDP-xylose does not. While *UXS1* orthologs are not found in *C. albicans* or *S. cerevisiae*, which lack xylose modifications, there is a *UXS1* ortholog in humans. 5FU is commonly used as a chemotherapeutic drug [40], and resistance to 5FU is frequently associated with mutations in thymidylate synthase [41]. Data here suggest that *uxs1* mutations may be acting in a similar fashion to either de-repress thymidylate synthase or inhibit Fur1 (Figure 5). Further exploration of the role of Uxs1 orthologs in humans during 5FU chemotherapy may be of interest.

## Material and methods

### Strains and media

The strains and plasmids used in this study are listed in Table S1. The strains were maintained in 25% glycerol stocks at −80°C and grown on rich YPD media at 30°C (Yeast extract Peptone Dextrose). Strains with selectable markers were grown on YPD containing 100 µg/mL nourseothricin (NAT) and/or 200 µg/mL G418 (NEO).

### Genome sequencing

DNA was isolated for sequencing by expanding individual colonies to 50 mL liquid cultures in YPD at 30°C. Cultures were then frozen and lyophilized until dry. DNA was extracted using a standard CTAB extraction protocol as previously described [42]. Illumina paired-end libraries were prepared and sequenced by the University of North Carolina Next Generation Sequencing Facility using the Kapa Library prep kit and the Hiseq platform. Additional sequencing was performed by the Duke University Sequencing Core using the Kapa Hyper prep kit and performed using a NovaSeq platform. Raw reads are available through the Sequence Read Archive under project accession number PRJNA525019.

### Genome assembly and variant calling

Reads were aligned to the V2 R265 reference genome [43] using BWA-MEM [44]. Alignments were further processed with SAMtools [45], the Genome Analysis Toolkit (GATK) [46], and Picard. SNP and indel calling was performed using the Unified Genotyper Component of the GATK with default settings aside from ploidy=1. VCFtools [47] was utilized for processing of the resulting calls to remove sites common to all strains (errors in the reference assembly) and variants were annotated using SnpEff [27]. All remaining variant calls were visually examined using the Integrated Genome Viewer (IGV) to remove calls resulting from poor read mapping [48]. FungiDB was also used to determine putative function and orthology of genes containing called variants in the dataset [49].

### Strain construction

A *ugd1*Δ mutant was constructed in the KN99**a** background as follows. Primers pairs JOHE45233/JOHE45085, JOHE45086/JOHE45087, and JOHE45088/JOHE45234 were used to amplify 1 kb upstream of *UGD1*, the neomycin resistant marker, and 1 kb downstream of the *UGD1* gene, respectively (primer sequences available in Table S2). To generate the deletion allele for *C. neoformans* transformation, all three fragments were cloned into plasmid pRS426 by transforming *S. cerevisiae* strain FY834 as previously described [50]. Recombinant *S. cerevisiae* transformants were selected on SD-uracil media and verified by spanning PCR with primer pair JOHE45233/JOHE45234. The resulting PCR product was introduced into *C. neoformans* laboratory strain KN99**a** by biolistic transformation and transformants were selected on YPD containing neomycin. Putative *ugd1*Δ deletion mutants were confirmed by PCR.

*uxs1*Δ single mutants and *ugd1*Δ *uxs1*Δ double mutants were generated via a genetic cross [51]. First, the KN99α *uxs1*Δ mutant from the Hiten Madhani deletion collection was mated with the wild-type KN99**a** laboratory strain. Through microdissection, spores were isolated, germinated, and genotyped via PCR for the gene deletion and the mating type locus to isolate a *MAT***a** *uxs1*Δ mutant in the KN99 background. Second, the KN99**a** *uxs1*Δ mutant was mated with wild-type H99. Spores were dissected and genotyped via PCR for the gene deletion and the mating type locus to isolate H99 *uxs1*Δ single mutants. Finally, the H99 *uxs1*Δ single mutant was crossed with KN99**a** *ugd1*Δ to generate *ugd1*Δ *uxs1*Δ double mutants, and the H99 *ugd1*Δ single mutant.

### Spot dilution assays

Single colonies were inoculated into 5 mL of liquid YPD and grown overnight at 30°C. Cell density was determined using a hemocytometer and the cultures were diluted accordingly such that 100,000 cells were aliquoted on to the most concentrated spot and subsequent spots consisted of 10-fold dilutions per spot. Each strain was spotted onto YPD or YNB alone and onto media also containing 5FC or 5FU at the indicated concentration. Plates were incubated at 30°C until photographed.

### Swab assays

Swab assays were conducted as previously described [13]. To isolate independent drug resistant strains, the original parent strains were subcultured from a frozen glycerol stock. Single colonies were used to inoculate liquid YPD cultures without selection. Those liquid cultures were grown with shaking until saturation. They were then spread onto drug plates (100 µg/mL 5FC on YNB) using sterile cotton swabs to select for resistant colonies. A single drug resistant colony was taken from any given liquid culture to ensure independence. This assay is only semi-quantitative, as the inoculum is not strictly controlled between independent cultures when swabbing.

### MIC Testing

5-flucytosine was dissolved in water and added to liquid YPD media in a 96-well plate at 400 µg/mL. 2-fold serial dilutions were performed until a concentration of 1.56 µg/mL was achieved. For YNB, 5-flucytosine was dissolved in water and added to liquid YNB media in a 96-well plate at 4 µg/mL. 2-fold serial dilutions were performed until a concentration of 0.016 µg/mL was achieved. Cell density of overnight cultures (liquid YPD, 30°C) was determined using a hemocytometer and cultures were adjusted to 10^5^ cells/mL. 100 µl of cell suspension was added to each well (10,000 cells per well). The 96-well plate was incubated at 30°C and OD600 readings were taken daily using a Sunrise Tecan instrument and Magellan software.

## Data Availability

Raw reads are available through the Sequence Read Archive under project accession number PRJNA525019. Strains generated in this study are available upon request.

## Acknowledgements

This study was supported by NIH/NIAID R37 MERIT award AI39115-21, NIH/NIAID R01 AI50113-15, NIH/NIAID R01 AI112595-04, and NIH/NIAID P01 AI104533-05 to J.H.; NIH/NIAID R21 AI109623 to T.L.D; and NIH/NIAID F30 AI120339 to L.X.L. This study utilized a *Cryptococcus* gene deletion collection deposited at the Fungal Genetics Stock Center and made freely available ahead of publication by the Madhani laboratory and funded by NIH R01 AI100272.

## Figure legends

**Supplementary Figure 1. VGIIa-like isolates acquire resistance to 5FC and 5FU more rapidly than the VGIIa isolate R265.**

VGIIa-like strains NIH444 and CBS7750 that harbor *msh2* nonsense alleles were tested for the ability to generate resistance to 5FC and 5FU in comparison with the closely related VGIIa strain R265. For each strain, 5 mL YPD cultures were inoculated from a single colony and grown overnight at 30°C. After washing, 100 µl of a 10^−5^ dilution was plated to YNB control plates and 100 µl of undiluted cultures was plated on media containing 5FC or 5FU. The VGIIa-like strains generated substantially more isolates resistant to both drugs.

**Supplementary Figure 2. Mutants of *fcy1* and *fur1* in *Cryptococcus neoformans* are resistant to 5FC but not 5FU.**

*fur1*Δ and *fcy1*Δ strains from the KN99 *C. neoformans* collection were struck onto YNB, YNB + 100 µg/mL 5FC, and YNB + 100 µg/mL 5FU. While the *fcy1*Δ mutant strain grew on media containing 5FC, it did not grow on media containing 5FU. In contrast, the *fur1*Δ mutant strain grew on media with either drug.

**Table S1.**
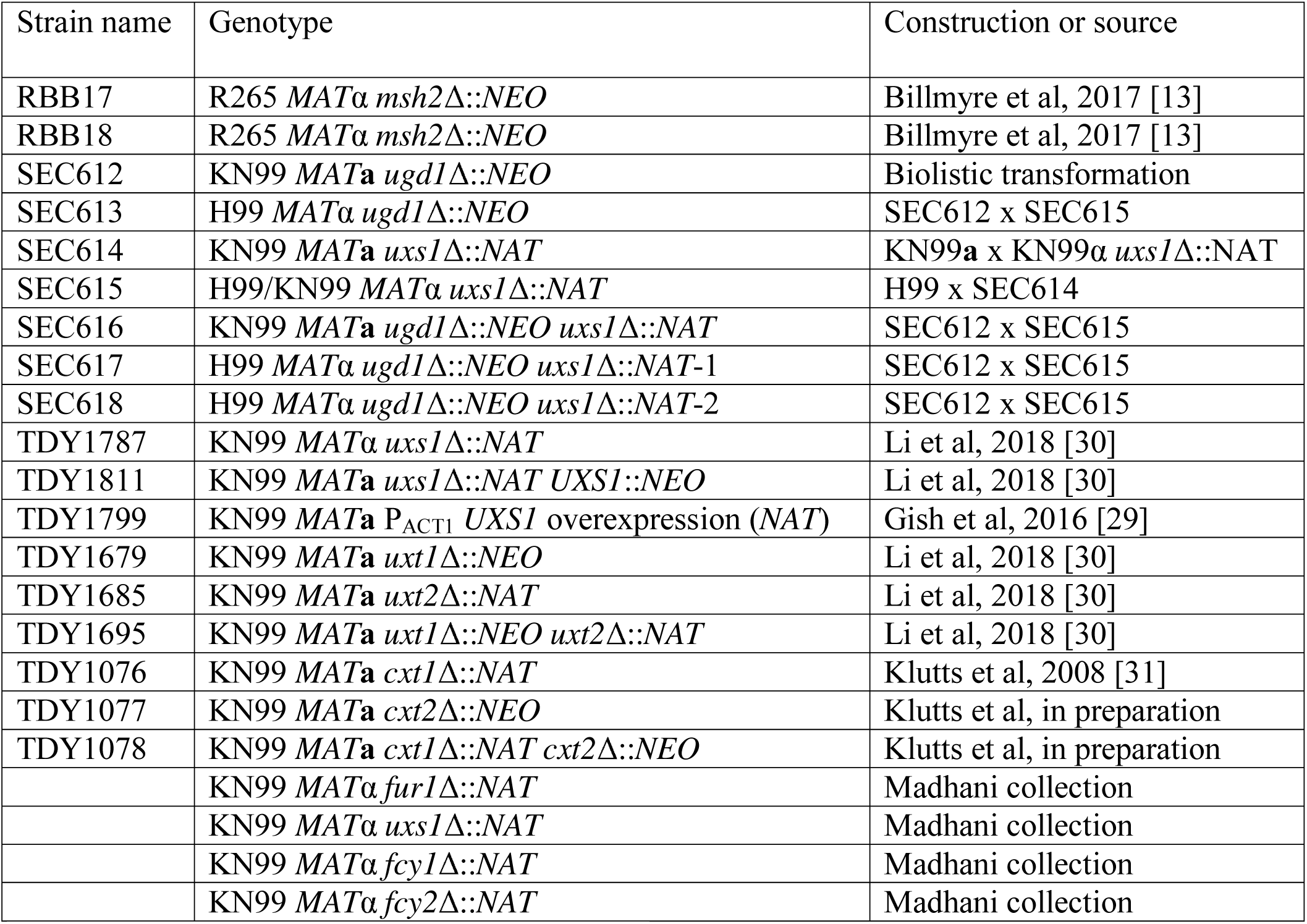
Strains and plasmids used in this study.

**Table S2.**
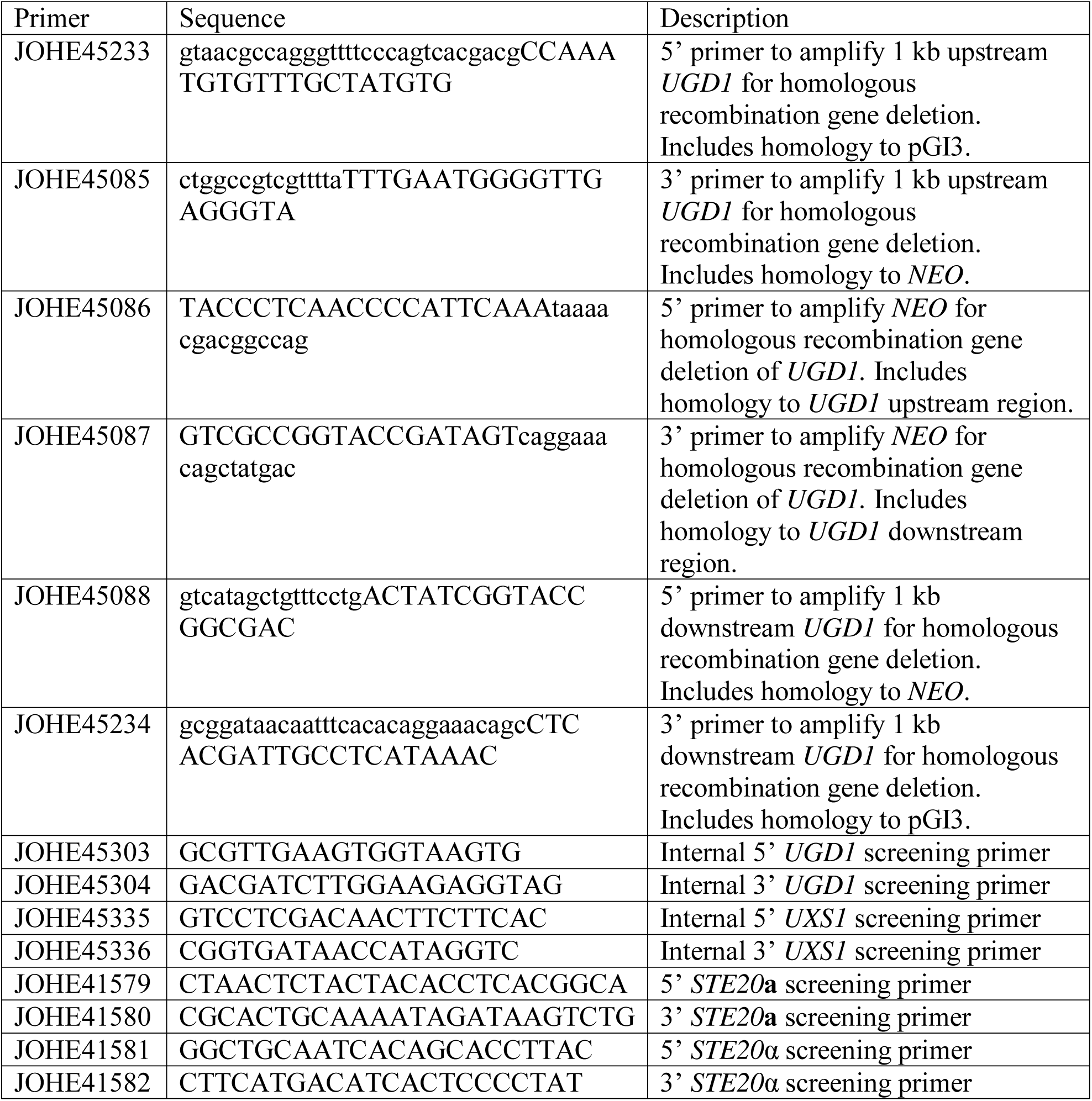
Oligonucleotides used in this study.

## References

1. Brown GD, Denning DW, Gow NAR, Levitz SM, Netea MG, White TC. Hidden killers: human fungal infections. Sci Transl Med. 2012;4: 165rv13.

2. Rajasingham R, Smith RM, Park BJ, Jarvis JN, Govender NP, Chiller TM, et al. Global burden of disease of HIV-associated cryptococcal meningitis: an updated analysis. Lancet Infect Dis. 2017;17: 873–881.

3. Saag MS, Graybill RJ, Larsen RA, Pappas PG, Perfect JR, Powderly WG, et al. Practice guidelines for the management of Cryptococcal disease. Clin Infect Dis. 2000;30: 710–718.

4. Bennett JE, Dismukes WE, Duma RJ, Medoff G, Sande MA, Gallis H, et al. A comparison of amphotericin B alone and combined with flucytosine in the treatment of cryptoccal meningitis. N Engl J Med. 1979;301: 126–131.

5. Molloy SF, Kanyama C, Heyderman RS, Loyse A, Kouanfack C, Chanda D, et al. Antifungal combinations for treatment of Cryptococcal meningitis in Africa. N Engl J Med. 2018;378: 1004–1017.

6. Loyse A, Dromer F, Day J, Lortholary O, Harrison TS. Flucytosine and cryptococcosis: time to urgently address the worldwide accessibility of a 50-year-old antifungal. J Antimicrob Chemother. 2013;68: 2435–2444.

7. Hope W, Tabernero L, Denning D, Anderson M. Molecular mechanisms of primary resistance to flucytosine in *Candida albicans*. Antimicrob Agents Chemother. 2004;48: 4377–4386.

8. Papon N, Noël T, Florent M, Gibot-Leclerc S, Jean D, Chastin C, et al. Molecular mechanism of flucytosine resistance in *Candida lusitaniae*: contribution of the *FCY2, FCY1*, and *FUR1* genes to 5-fluorouracil and fluconazole cross-resistance. Antimicrob Agents Chemother. 2007;51: 369–371.

9. McManus BA, Moran GP, Higgins JA, Sullivan DJ, Coleman DC. A Ser29Leu substitution in the cytosine deaminase Fca1p is responsible for clade-specific flucytosine resistance in *Candida dubliniensis*. Antimicrob Agents Chemother. 2009;53: 4678–4685.

10. Whelan WL. The genetic basis of resistance to 5-fluorocytosine in *Candida* species and *Cryptococcus neoformans*. CRC Crit Rev Microbiol. 1987;15: 45–56.

11. Vu K, Thompson George R III, Roe CC, Sykes JE, Dreibe EM, Lockhart SR, et al. Flucytosine resistance in *Cryptococcus gattii* is indirectly mediated by the *FCY2-FCY1-FUR1* pathway. Med Mycol. 2018;56: 857–867.

12. Khanal Lamichhane A, Garraffo HM, Cai H, Walter PJ, Kwon-Chung KJ, Chang YC. A novel role of fungal type I myosin in regulating membrane properties and its association with D-amino acid utilization in *Cryptococcus gattii*. mBio. 2019;10: e01867–19.

13. Billmyre RB, Clancey SA, Heitman J. Natural mismatch repair mutations mediate phenotypic diversity and drug resistance in *Cryptococcus deuterogattii*. eLife. 2017;6: e28802.

14. Boyce KJ, Wang Y, Verma S, Shakya VPS, Xue C, Idnurm A. Mismatch repair of DNA replication errors contributes to microevolution in the pathogenic fungus *Cryptococcus neoformans*. mBio. 2017;8.

15. Billmyre RB, Croll D, Li W, Mieczkowski P, Carter DA, Cuomo CA, et al. Highly recombinant VGII *Cryptococcus gattii* population develops clonal outbreak clusters through both sexual macroevolution and asexual microevolution. mBio. 2014;5: e01494–14.

16. Rhodes J, Beale MA, Vanhove M, Jarvis JN, Kannambath S, Simpson JA, et al. A population genomics approach to assessing the genetic basis of within-host microevolution underlying recurrent cryptococcal meningitis infection. G3. 2017;7: 1165–1176.

17. Healey KR, Zhao Y, Perez WB, Lockhart SR, Sobel JD, Farmakiotis D, et al. Prevalent mutator genotype identified in fungal pathogen *Candida glabrata* promotes multi-drug resistance. Nat Commun. 2016;7: 11128.

18. Dellière S, Healey K, Gits-Muselli M, Carrara B, Barbaro A, Guigue N, et al. Fluconazole and echinocandin resistance of *Candida glabrata* correlates better with antifungal drug exposure rather than with *MSH2* mutator genotype in a French cohort of patients harboring low rates of resistance. Front Microbiol. 2016;7: 2038.

19. Healey KR, Jimenez Ortigosa C, Shor E, Perlin DS. Genetic drivers of multidrug resistance in *Candida glabrata*. Front Microbiol. 2016;7: 1995.

20. Singh A, Healey KR, Yadav P, Upadhyaya G, Sachdeva N, Sarma S, et al. Absence of azole or echinocandin resistance in *Candida glabrata* isolates in India despite background prevalence of strains with defects in the DNA mismatch repair pathway. Antimicrob Agents Chemother. 2018;62: e00195–18.

21. Steenwyk JL, Opulente DA, Kominek J, Shen X-X, Zhou X, Labella AL, et al. Extensive loss of cell-cycle and DNA repair genes in an ancient lineage of bipolar budding yeasts. PLOS Biol. 2019;17: e3000255.

22. Hagen F, Khayhan K, Theelen B, Kolecka A, Polacheck I, Sionov E, et al. Recognition of seven species in the *Cryptococcus gattii*/*Cryptococcus neoformans* species complex. Fungal Genet Biol. 2015;78: 16–48.

23. Rhodes J, Desjardins CA, Sykes SM, Beale MA, Vanhove M, Sakthikumar S, et al. Tracing genetic exchange and biogeography of *Cryptococcus neoformans* var. *grubii* at the global population level. Genetics. 2017;207: 327–346.

24. Kwon-Chung KJ, Bennett JE, Wickes BL, Meyer W, Cuomo CA, Wollenburg KR, et al. The case for adopting the “Species Complex” nomenclature for the etiologic agents of Cryptococcosis. mSphere. 2017;2.

25. Gsaller F, Furukawa T, Carr PD, Rash B, Jöchl C, Bertuzzi M, et al. Mechanistic basis of pH-dependent 5-flucytosine resistance in *Aspergillus fumigatus*. Antimicrob Agents Chemother. 2018;62: e02593–17.

26. Yadav V, Sun S, Billmyre RB, Thimmappa BC, Shea T, Lintner R, et al. RNAi is a critical determinant of centromere evolution in closely related fungi. Proc Natl Acad Sci. 2018;115: 3108–3113.

27. Cingolani P, Platts A, Wang LL, Coon M, Nguyen T, Wang L, et al. A program for annotating and predicting the effects of single nucleotide polymorphisms, SnpEff: SNPs in the genome of *Drosophila melanogaster* strain w1118; iso-2; iso-3. Fly. 2012;6: 80–92.

28. Bar-Peled M, Griffith CL, Doering TL. Functional cloning and characterization of a UDP-glucuronic acid decarboxylase: the pathogenic fungus *Cryptococcus neoformans* elucidates UDP-xylose synthesis. Proc Natl Acad Sci. 2001;98: 12003 LP – 12008.

29. Gish SR, Maier EJ, Haynes BC, Santiago-Tirado FH, Srikanta DL, Ma CZ, et al. Computational analysis reveals a key regulator of cryptococcal virulence and determinant of host response. mBio. 2016;7: e00313–16.

30. Li LX, Rautengarten C, Heazlewood JL, Doering TL. Xylose donor transport is critical for fungal virulence. PLoS Pathog. 2018;14: e1006765.

31. Klutts JS, Doering TL. Cryptococcal xylosyltransferase 1 (Cxt1p) from *Cryptococcus neoformans* plays a direct role in the synthesis of capsule polysaccharides. J Biol Chem. 2008;283: 14327–14334.

32. Griffith CL, Klutts JS, Zhang L, Levery SB, Doering TL. UDP-glucose dehydrogenase plays multiple roles in the biology of the pathogenic fungus *Cryptococcus neoformans*. J Biol Chem. 2004;279: 51669–51676.

33. Harris BE, Manning BW, Federle TW, Diasio RB. Conversion of 5-fluorocytosine to 5-fluorouracil by human intestinal microflora. Antimicrob Agents Chemother. 1986;29: 44–48.

34. Tenaillon O, Le Nagard H, Godelle B, Taddei F. Mutators and sex in bacteria: conflict between adaptive strategies. Proc Natl Acad Sci U S A. 2000;97: 10465–70.

35. Moyrand F, Klaproth B, Himmelreich U, Dromer F, Janbon G. Isolation and characterization of capsule structure mutant strains of *Cryptococcus neoformans*. Mol Microbiol. 2002;45: 837–849.

36. Leipheimer J, Bloom ALM, Baumstark T, Panepinto JC. CNBP homologues Gis2 and Znf9 interact with a putative G-quadruplex-forming 3’ untranslated region, altering polysome association and stress tolerance in *Cryptococcus neoformans*. mSphere. 2018;3: e00201–18.

37. Schwarz P, Janbon G, Dromer F, Lortholary O, Dannaoui E. Combination of amphotericin B with flucytosine is active in vitro against flucytosine-resistant isolates of *Cryptococcus neoformans*. Antimicrob Agents Chemother. 2007;51: 383–385.

38. Chapeland-Leclerc F, Bouchoux J, Goumar A, Chastin C, Villard J, Noel T. Inactivation of the *FCY2* gene encoding purine-cytosine permease promotes cross-resistance to flucytosine and fluconazole in *Candida lusitaniae*. Antimicrob Agents Chemother. 2005;49: 3101–3108.

39. Florent M, Noel T, Ruprich-Robert G, Da Silva B, Fitton-Ouhabi V, Chastin C, et al. Nonsense and missense mutations in *FCY2* and *FCY1* genes are responsible for flucytosine resistance and flucytosine-fluconazole cross-resistance in clinical isolates of *Candida lusitaniae*. Antimicrob Agents Chemother. 2009;53: 2982–2990.

40. Moertel CG. Chemotherapy for colorectal cancer. N Engl J Med. 1994;330: 1136–1142.

41. Pullarkat ST, Stoehlmacher J, Ghaderi V, Xiong Y-P, Ingles SA, Sherrod A, et al. Thymidylate synthase gene polymorphism determines response and toxicity of 5-FU chemotherapy. Pharmacogenomics J. 2001;1: 65.

42. Pitkin JW, Panaccione DG, Walton JD. A putative cyclic peptide efflux pump encoded by the *TOXA* gene of the plant-pathogenic fungus *Cochliobolus carbonum*. Microbiology. 1996;142: 1557–1565.

43. Farrer RA, Desjardins CA, Sakthikumar S, Gujja S, Saif S, Zeng Q, et al. Genome evolution and innovation across the four major lineages of *Cryptococcus gattii*. mBio. 2015;6: e00868–15.

44. Li H, Durbin R. Fast and accurate short read alignment with Burrows-Wheeler transform. Bioinformatics. 2009;25: 1754–60.

45. Li H, Handsaker B, Wysoker A, Fennell T, Ruan J, Homer N, et al. The Sequence Alignment/Map format and SAMtools. Bioinformatics. 2009;25: 2078–9.

46. McKenna A, Hanna M, Banks E, Sivachenko A, Cibulskis K, Kernytsky A, et al. The Genome Analysis Toolkit: A MapReduce framework for analyzing next-generation DNA sequencing data. Genome Res. 2010;20: 1297–303.

47. Danecek P, Auton A, Abecasis G, Albers CA, Banks E, DePristo MA, et al. The variant call format and VCFtools. Bioinformatics. 2011;27: 2156–8.

48. Thorvaldsdóttir H, Robinson JT, Mesirov JP. Integrative Genomics Viewer (IGV): high-performance genomics data visualization and exploration. Brief Bioinform. 2013;14: 178–92.

49. Stajich JE, Harris T, Brunk BP, Brestelli J, Fischer S, Harb OS, et al. FungiDB: an integrated functional genomics database for fungi. Nucleic Acids Res. 2012;40: D675–81.

50. Ianiri G, Averette AF, Kingsbury JM, Heitman J, Idnurm A. Gene function analysis in the ubiquitous human commensal and pathogen *Malassezia genus*. mBio. 2016;7.

51. Sun S, Priest SJ, Heitman J. *Cryptococcus neoformans* mating and genetic crosses. Curr Protoc Microbiol. 2019;53: e75.

