## Supplementary figures and images for "Hypermutation in *Cryptococcus* reveals a novel pathway to 5-fluorocytosine (5FC) resistance"

### Supplemental Figure 1

#### Supplemental Figure 1

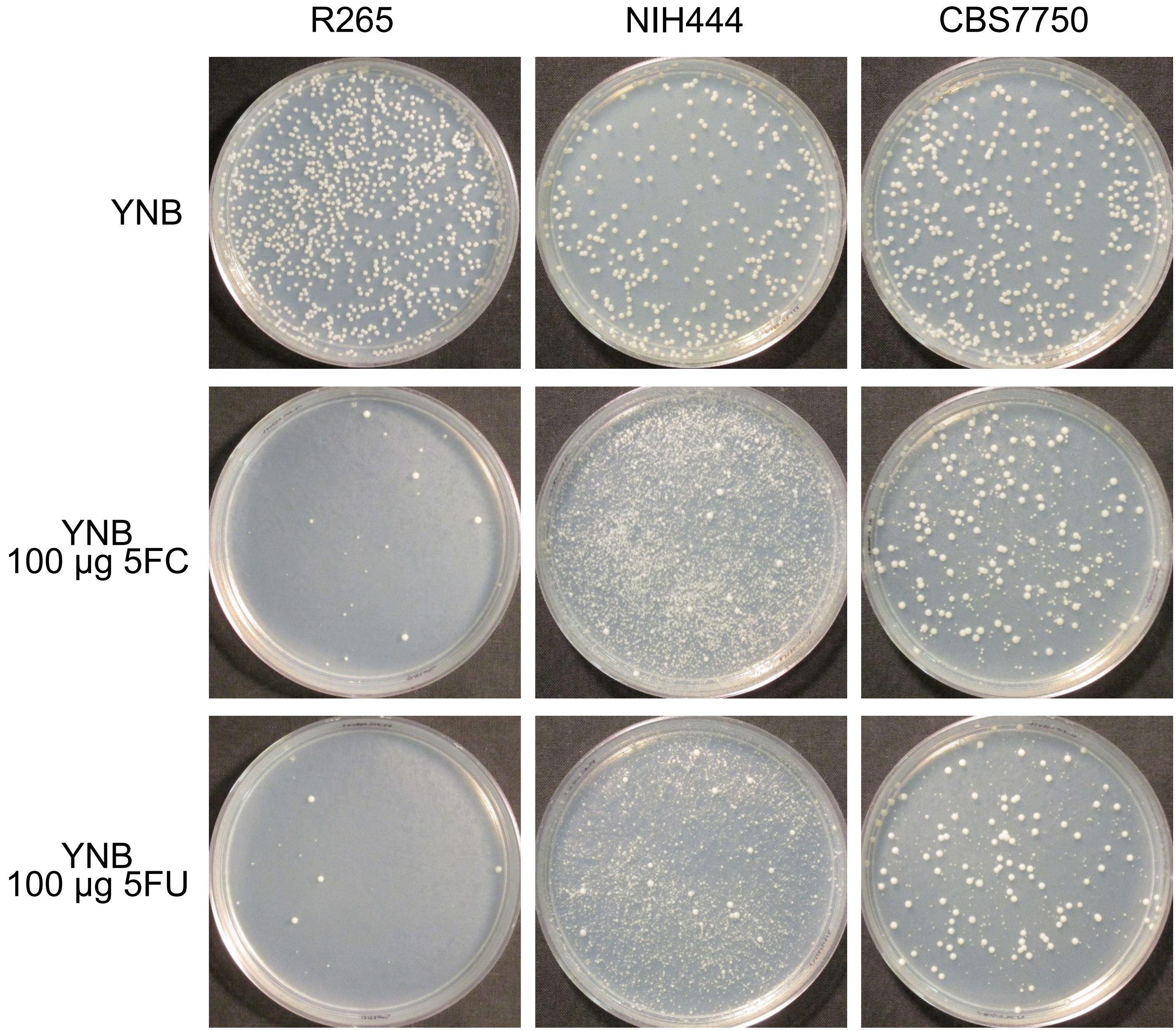

### Supplemental Figure 2

**Supplemental Figure 2**

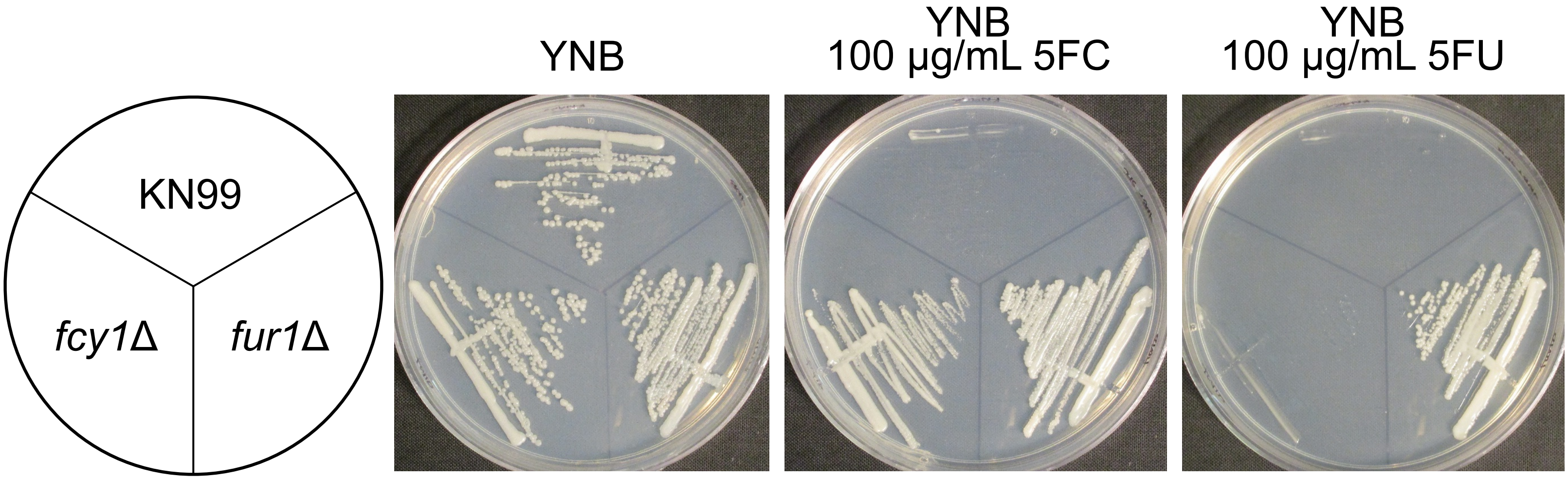
